# Differential small RNA responses against co-infecting insect-specific viruses in *Aedes albopictus* mosquitoes

**DOI:** 10.1101/2020.03.28.012062

**Authors:** Lionel Frangeul, Hervé Blanc, Maria-Carla Saleh, Yasutsugu Suzuki

**Affiliations:** Institut Pasteur, Viruses and RNA Interference Unit, CNRS Unité Mixte de Recherche 3569, 75724 Paris Cedex 15, France

**Keywords:** *Aedes albopictus*, insect-specific viruses, co-infection, small interfering RNA, PIWI-interacting RNA, reproductive tissues, sex difference

## Abstract

The mosquito antiviral response has been mainly studied in the case of arthropod-borne virus (arbovirus) infection in female mosquitoes. However, in nature, both female and male mosquitoes are abundantly infected with insect-specific viruses (ISVs). ISVs are capable of infecting the reproductive organs of both sexes and are maintained primarily by vertical transmission. Since the RNA interference (RNAi)-mediated antiviral response plays an important antiviral role in mosquitoes, ISVs constitute a relevant model to study sex-dependent antiviral responses. Using a naturally generated viral stock containing three distinct ISVs, *Aedes* flavivirus (AEFV), Menghai Rhabdovirus (MERV) and Shinobi tetra virus (SHTV), we infected adult *Aedes albopictus* females and males and generated small RNA libraries from ovaries, testes, and the remainder of the body. Overall, both female and male mosquitoes showed unique small RNA profiles to each co-infecting ISV regardless the sex or tissue tested. While all three ISVs generated virus-derived siRNAs, only MERV generated virus-derived piRNAs. We also studied the expression of PIWI genes in reproductive tissues and carcasses. Piwi1-4 were abundantly expressed in ovaries and testes in contrast to Piwi5-9, suggesting that Piwi 5-9 are involved in exogenous viral piRNA production. Together, our results show that ISV-infected *Aedes albopictus* produce viral small RNAs in a virus-specific manner and that male mosquitoes mount a similar small RNA-mediated antiviral response to that of females.

## 1. Introduction

Mosquitoes are efficient vectors for various human pathogens. *Aedes* and *Culex* are two major genera of mosquitoes that transmit medically important arthropod-borne viruses (arboviruses) such as dengue, West Nile and Zika viruses. Recent metagenomics studies have shed light on the virome of these mosquito vectors, revealing diverse virus communities largely composed of insect-specific viruses (ISVs) that specifically infect invertebrates but not vertebrates [1-7]. ISVs have been shown to negatively regulate some arbovirus infections in mosquito cell cultures and *in vivo* [8-14]. For instance, two ISVs, Menghai rhabdovirus (MERV) and Shinobi tetravirus (SHTV), which were identified in an *Aedes* (*Ae*.) *albopictus* C6/36 cell line from the Japanese Collection of Research Bioresources (JCRB), suppressed Zika virus replication *in vitro* [14]. In spite of increased attention to ISVs, little is known about mosquito-ISV interactions, from the natural infection routes *in vivo* to the immune responses to ISVs. For example, although some ISVs induce cytopathic effects in mosquito cell cultures, few studies have investigated the *in vivo* pathogenesis. It has been suggested that mosquitoes establish a tolerant state to ISVs as well as to arboviruses and that this viral tolerance is one of the critical factors influencing mosquito vector competence [15-17]. Therefore, understanding the mosquito antiviral response to ISVs could provide new insights into the viral tolerance mechanisms of mosquitoes.

The small interfering RNA (siRNA) pathway is considered as a major antiviral defense machinery in insects including mosquito vectors [18]. The cellular RNase-III Dicer-2 (DCR2) recognizes viral dsRNA forms, which are mostly viral replication intermediates and viral secondary structures, and processes them into 21 nt-long siRNAs. The siRNA duplexes are loaded into the RNA-induced silencing complex (RISC) containing Argonaute2 (Ago2) that cleaves the target viral RNA under the guide of siRNA. Some mosquito viruses have evolved avoidance mechanisms to the siRNA pathway, demonstrating that virus-derived siRNAs (vsiRNAs) play an important antiviral function in mosquito vectors [19-24].

The P-element induced wimpy testis (PIWI)-interacting RNA (piRNA) pathway has been demonstrated to control transposable elements (TEs) in germ lines. Biogenesis of piRNAs is well characterized in *Drosophila melanogaster* and reviewed in [25]. piRNAs can be divided into two different classes: primary piRNAs and secondary piRNAs. Both types of piRNAs show a similar size of 25-30 nt. Primary piRNAs are generated from single-stranded precursor transcripts from piRNA clusters enriched in transposons remnants, and their first nucleotide is typically a uridine, referred to as 1U bias. Secondary piRNAs are produced by cleavage events of complementary transcripts by primary piRNAs and often contain an adenine at the 10th nucleotide position, called 10A bias. In addition to piRNA cluster-derived piRNAs, mosquitoes have been shown to produce viral piRNAs (vpiRNAs) from different replicating viruses such as flaviviruses, alphaviruses and rhabdoviruses [15, 26-32]. This contrasts with *Drosophila melanogaster*, where vpiRNA are not produced during virus infections [33]. The process leading to the production of vpiRNAs remains unknown, mosquitoes having an expanded PIWI family compared to *Drosophila*; *Ae. aegypti* has eight members of PIWI genes (Piwi1-7 and Ago3) and *Ae. albopictus* was estimated to have ten PIWI genes (Piwi1-9 and Ago3) [34, 35]. This expansion could explain the production of vpiRNAs in mosquitoes. The antiviral potential of vpiRNAs remains unclear and the function of PIWI proteins in *Ae. albopictus* remains to be determined. Functional studies of PIWI proteins have been primarily conducted in the *Ae. aegypti* cell line Aag2 using arboviruses. Gene silencing of Piwi5 and Ago3 decreased vpiRNA biogenesis without impacting replication of Semliki Forest virus (SFV), Sindbis virus (SINV) and dengue virus [26, 27, 29, 31, 36]. Further experiments suggested that Piwi4 was antiviral against SFV, but not through the piRNA pathway [27]. More recently, it has been suggested that Piwi4 is involved in maturation of both siRNAs and piRNAs [37].

Unlike arboviruses, ISVs are likely to be commensal in mosquito hosts and are considered to be vertically transmitted to the offspring for their maintenance in nature [38-43]. Due to their mode of transmission, ISVs provide a great opportunity to better understand the mosquito antiviral response in females as well as in males, especially the antiviral responses such as the piRNA pathway in reproductive organs and germ line. To date, studies on ISV-derived small RNA production have been limited to mosquito cell lines [28, 30]. Here, we investigated the small RNA responses to ISVs in individual mosquitoes. A previous study identified *Aedes* flavivirus (AEFV) in a field-collected *Ae. albopictus* individual [44]. The AEFV strain was isolated in the C6/36 cell line from JCRB, which is persistently infected with two other ISVs, MERV and SHTV [44]. Consequently, AEFV stocks generated in these cells contain three ISVs that are phylogenetically distant: AEFV belongs to the *Flaviviridae* family and is a positive-sense single-stranded RNA (ssRNA) virus; MERV is from the *Rhabdoviridae* and is negative-sense ssRNA virus; and SHTV is from the *Tetraviridae* and is a positive-sense ssRNA virus. To compare the small RNA responses generated by these distant ISVs, we deep-sequenced small RNAs from ovaries, testes and carcasses of *Ae. albopictus* triple-infected with AEFV, MERV and SHTV. All three viruses infected the reproductive tissues and carcasses. Each ISV showed a unique small RNA profile independent of the sex and tissue of the mosquito tested. The testes produced comparable amounts of viral small RNAs as the carcasses while the ovaries produced reduced amounts. We also observed differential gene expression for some PIWI-genes between reproductive tissues and the carcasses. Our results showed that *Ae. albopictus* responds to co-infecting ISVs in a virus-specific manner at the small RNA level and that male adult mosquitoes mount a similar small RNA response to that of females during viral infection.

## 2. Materials and Methods

### 2. 1. Cell culture and virus production

C6/36 cells (ATCC CRL-1660) derived from *Ae. albopictus* were maintained at 28°C in L-15 Leibovitz’s medium (Gibco) supplemented with 10% fetal bovine serum (FBS; Gibco), 1% nonessential amino acids (Gibco), 2% tryptose phosphate broth (Sigma), and 1% penicillin-streptomycin (Gibco). The cells were tested by next-generation sequencing (NGS) and are free of known viruses present in the NCBI viruses database.

AEFV strain Narita-21/MERV isolate Menghai/SHTV strain Shinobi mixture stock was kindly provided by Haruhiko Isawa, National Institute of Infectious Diseases, Japan. The virus stock was amplified in naïve C6/36 cells grown in L-15 Leibovitz’s medium and the supernatant was used for the co-infection experiment *in vivo*. To create a standard curve for AEFV, the non-structural gene 4 (NS4) was amplified with primers containing the T7 promoter sequence followed by reverse transcription with MEGAscript T7 Transcription Kit (Thermo Fisher) to generate AEFV NS4 RNA. The copy number of AEFV RNA was determined by reverse transcription quantitative polymerase chain reaction (RT-qPCR) based on serial dilution of the AEFV NS4 RNA with Maxima H minus First Strand cDNA Synthesis kit (Thermo Fisher) and Fast SYBR Green Master Mix (Thermo Fisher). MERV and SHTV RNA copy numbers in the virus stock were calculated using Qgene template program [45].

### 2. 2. Mosquitoes and virus infections

Laboratory colonies of *Ae. albopictus* were established from field collections in Vietnam (2011) and in Japan (2009). The insectary conditions for maintenance of mosquitoes were 28°C, 70% relative humidity and a 12 h light: 12 h dark cycle. Adult mosquitoes were maintained with permanent access to 10% sucrose solution.

*Ae. albopictus* adult females and males, 4-7 days old, were intrathoracically injected with an AEFV/MERV/SHTV mixture containing 1 x 10^7^ genome copies of AEFV, 3.2 x 10^5^ genome copies of MERV, and 1.6 x 10^5^ genome copies of SHTV with a nanoinjector (Nanoject III, Drummond Scientific). Following the injection, mosquitoes were incubated at 28°C, 70% relative humidity and a 12 h light: 12 h dark cycle with permanent access to 10% sucrose solution.

### 2. 3. *Quantification of ISVs, siRNA and piRNA pathway-related gene RNA levels in co-infected* Ae. albopictus *mosquitoes*

Ovaries, testes and the rest of the bodies, which we define as carcasses for this study, from the AEFV/MERV/SHTV-injected mosquitoes were manually dissected at day 8 post infection. Total RNA was extracted from 3 pools of 20 ovaries, testes and carcasses each with TRIzol reagent (Invitrogen). cDNA synthesis and qPCR were performed as described above for AEFV. To normalize ISV, DCR2, Ago2, and PIWI gene expression, two different housekeeping genes were used, actin and ribosomal protein L18 (RPL18). All PCR primer sequences are listed in Table S1.

### 2. 4. Small RNA library preparation

Three pools of 20 ovaries, testes or carcasses each from the ISV-infected mosquitoes were combined for generation of small RNA libraries. 19-33 nt long RNAs were cut and extracted from a 15% acrylamide/bis-acrylamide (37.5:1), 7M urea gel and the purified RNAs were subjected to small RNA library preparation using NEBNext Multiplex Small RNA Library Prep (Illumina, New England Biolabs) with 3’ adaptor from Integrated DNA Technologies and in-house designed indexed primers. Libraries diluted to 4 nM were sequenced with NextSeq 500 High Output Kit v2 (75 cycles) on a NextSeq 500 (Illumina). Small RNA libraries have been submitted to the NCBI sequence read archive (SRA) under BioProject PRJNA587399.

### 2. 5. Bioinformatics analysis

The quality of the fastq files was assessed with FastQC software (www.bioinformatics.babraham.ac.uk/projects/fastqc/). Low-quality bases and adaptors were trimmed from each read using the Cutadapt program and only reads showing an acceptable quality (Phred score, 20) were retained. A second set of graphics was generated by the FastQC software using the trimmed fastq files. Reads were mapped to the genome sequences of AEFV strain Narita-21 (GenBank accession no. AB488408.1), MERV isolate Menghai (GenBank accession no. KX785335.1) and SHTV strain shinobi (GenBank accession no. LC270813.1) using the Bowtie 1 tool with the -v 1 (one mismatch between the read and its target) with default options for small RNAs. The Bowtie 1 tool generates results in Sequence Alignment/Map (SAM) format. All SAM files were analyzed using the SAMtools package to produce bam indexed files. Homemade R scripts with Rsamtools and Shortreads in Bioconductor software were used for analysis of the bam files to create graphs. The number of small RNA reads mapped on each virus were normalized by the total number of reads in each small RNA library. To survey MERV-like DNA sequences in the genome of the *Ae. albopictus* Vietnam strain, a BLASTn search against its genomic DNA library, deposited in SRA under BioProject accession number PRJNA587399, was performed.

## 3. Results

### 3.1. *AEFV, MERV and SHTV infection in co-injected* Ae. albopictus *mosquitoes*

*Ae. albopictus*-Vietnam (colony derived from a field collection in Vietnam) adult females and males were intrathoracically injected with a naturally derived viral stock containing a mixture of three ISVs. The relative genome copies for each virus injected in the mosquito was as follow: 1 x 10^7^ genome copies of AEFV, 3.2 x 10^5^ genome copies of MERV, and 1.6 x 10^5^ genome copies of SHTV. At 8 days post injection (dpi), ovaries, testes and their respective carcasses, which refer to the rest of body parts without ovaries or testes, were dissected and 3 pools of 20 tissues each were prepared to examine ISV infections. Each ISV RNA level was quantified by RT-qPCR normalized with actin or RPL18 RNA (Figure 1A and B). ISV RNA levels were very similar across normalization with either actin or RPL18. Both AEFV and MERV infected the reproductive tissues and their carcasses. SHTV infection was also observed at least in one of the pools in all conditions although the amount of SHTV RNA was one half to four logs lower than for AEFV and MERV. Two pools of testes showed undetected levels of SHTV. Only female carcasses allowed relatively high SHTV replication while AEFV and MERV RNA levels were comparable between both females and males carcasses. This is in agreement with a previous report showing that these ISVs do not compete with each other during infection in mosquito C6/36 cell line [14].

**Figure 1.**
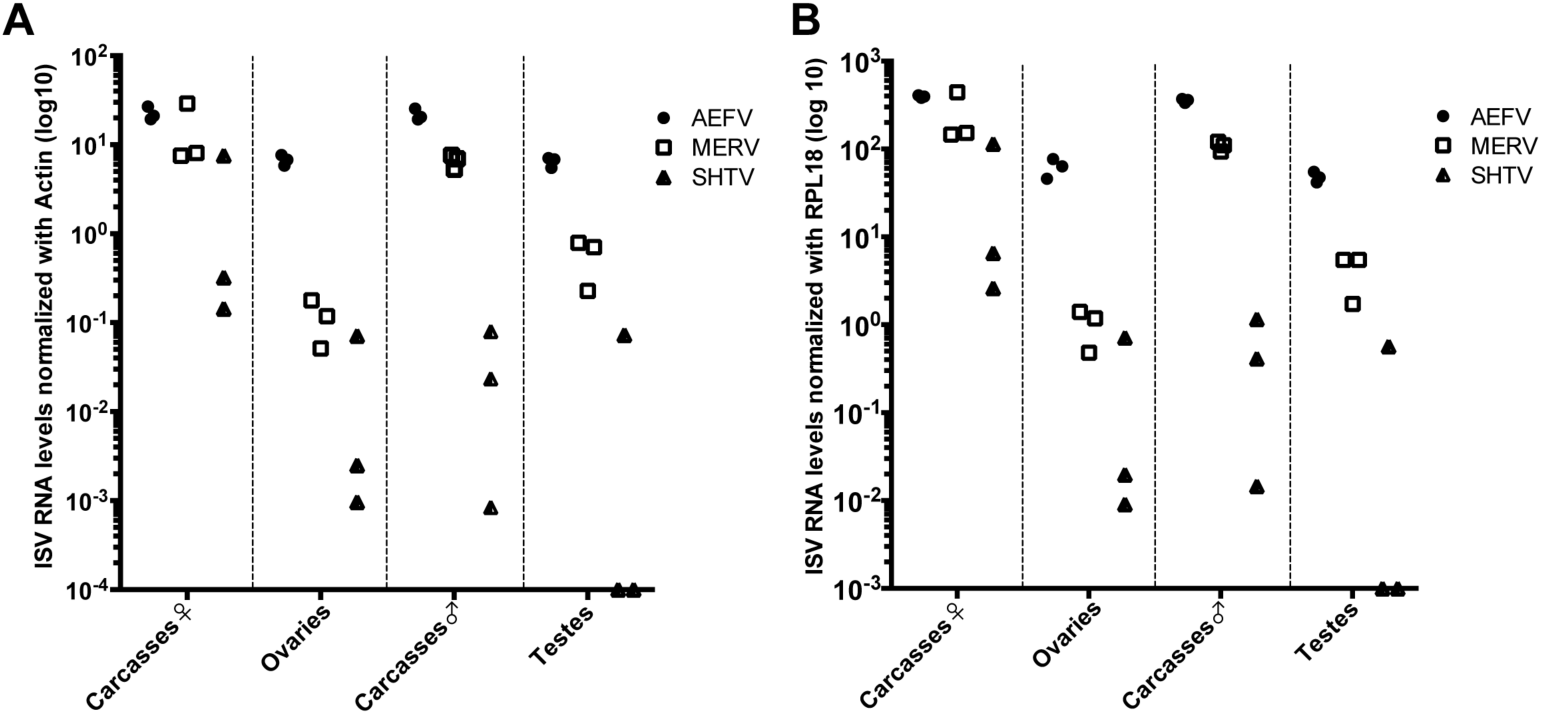
AEFV, MERV and SHTV infections in reproductive tissues (ovaries and testes) and carcasses. *Ae. albopictus* adult females and males were intrathoracically injected with an AEFV/MERV/SHTV mixture and collected at 8 days post injection. Each ISV RNA level in the coinfected mosquito ovaries, testes or their remaining carcasses was quantified by RT-qPCR normalized with actin (A) or RPL18 RNA (B). Each dot represents a pool of 20 reproductive tissues or carcasses.

### 3.2. *Small RNA profiles in AEFV, MERV and SHTV-co-infected* Ae. albopictus *mosquitoes*

In order to characterize small RNA responses to AEFV, MERV and SHTV in the co-infected *Ae. albopictus*-Vietnam mosquitoes, we generated small RNA libraries with the combined RNA samples from the 3 pools used for quantification of ISV RNA levels shown in Figure 1. The size distribution of the total small RNAs from both female and male carcasses showed peaks at 21 and 22 nt corresponding to siRNAs and miRNAs, respectively (Figure 2A, blue). Ovaries and testes showed an enrichment for 27-30 nt small RNAs corresponding to piRNAs (Figure 2A, red). Small RNA reads were mapped on each virus genome sequence to determine the virus-specific size distribution of virus-derived small RNAs. In both reproductive tissues and remaining carcasses, vsiRNAs of 21 nt were detected for each of the three viruses (Figure 2B and S1); and this was more pronounced in the carcasses of female mosquitoes. Ovaries produced approximately 0.39, 0.18 and 0.11% of total vsiRNAs from AEFV, MERV and SHTV, respectively, when compared to carcasses. Testes showed higher proportions of vsiRNAs from all three ISVs with 39%, 17% and 63% of total vsiRNAs from AEFV, MERV and SHTV respectively. A peak of 26-30 nt-long viral small RNAs, potentially vpiRNAs, was observed only for MERV (Figure S1). To compare putative vpiRNA production among the three ISVs, we calculated the proportion of each virus-derived 26-30 nt-long small RNAs to 100 reads of vsiRNAs targeting each virus respectively (Figure 2C). The proportion of MERV-derived putative piRNAs was between 30-90%, and similar between sexes and tissues tested. AEFV-derived putative piRNAs were detected at lower levels, 1-30% (Figure 2C and S1). However, unlike MERV-piRNAs, AEFV-piRNA-like small RNAs were preferably generated from the positive-strand orientation of the viral sequence, i.e. the viral genomes. SHTV showed almost no production of 26-30 nt-long small RNAs with a maximum of 2.5% compared to the vsiRNAs. To check that the absence of of 26-30 nt-long small RNAs from SHTV was not due to the lower level of viral replication (Figure 1 and 2B), we generated a new small RNA library from *Ae. albopictus-*Japan (colony derived from a field collection in Japan) co-infected with the three ISVs. The analysis of this library confirmed that, despite the high number of read corresponding to SHTV-derived siRNAs, carcasses of female mosquitoes showed almost an undetectable number of reads of 26-30 nt-long for this virus (Figure S2).

**Figure 2.**
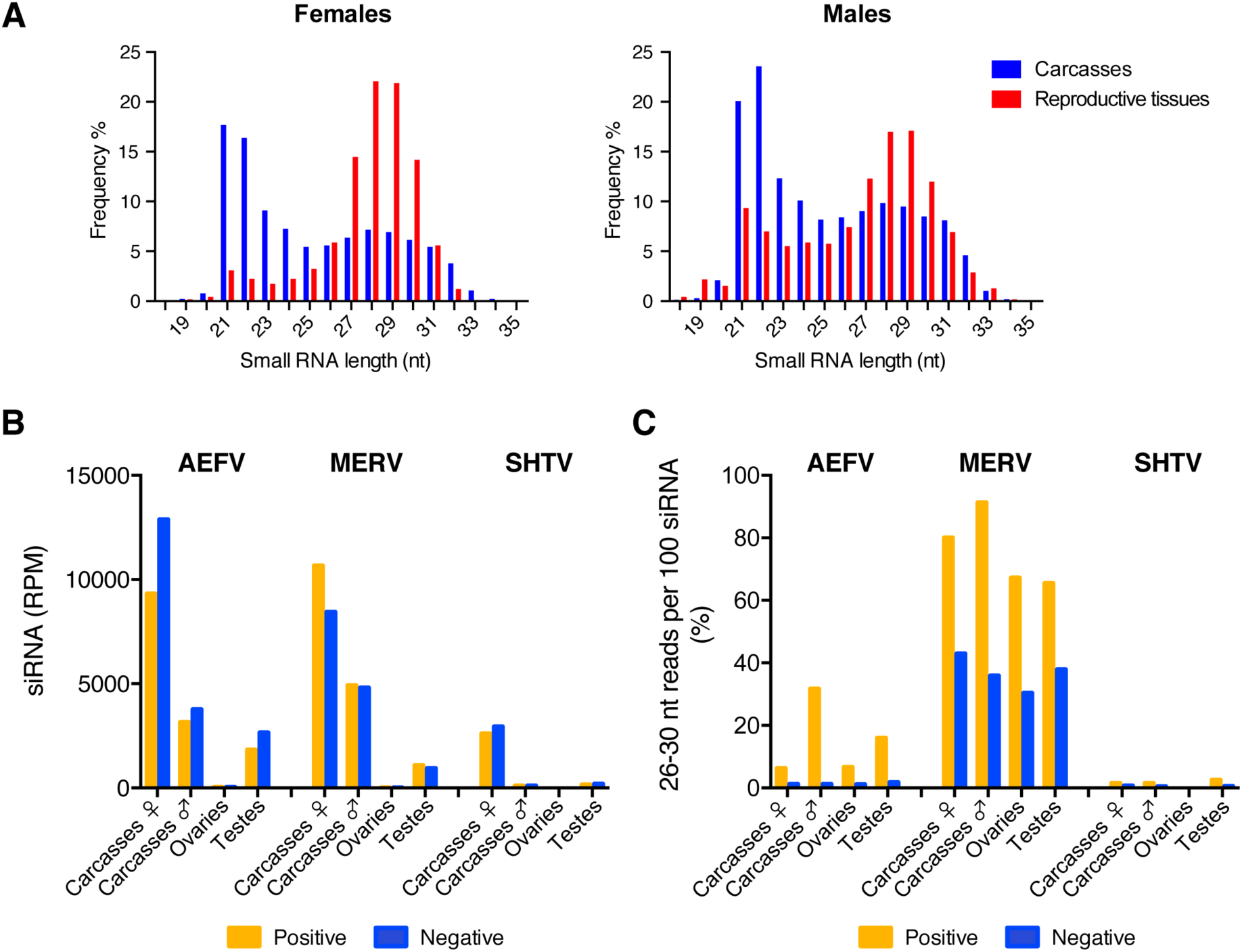
Comparison of vsiRNAs and putative vpiRNAs production from ISVs in reproductive tissues and carcasses of female or male *Ae. albopictus* mosquitoes. Frequency of the size distribution of total small RNA reads from each sample is shown in (A). Normalized siRNA reads per one million (RPM) mapped to AEFV, MERV or SHTV genome is shown in (B). Proportion of 26-30 nt long AEFV-, MERV- or SHTV-derived small RNAs to 100 reads of its vsiRNAs is represented in (C). Yellow and blue bars represent positive- and negative-stranded reads, respectively.

Overall, our results indicated that the type of virus-derived small RNAs produced during virus infection was virus-dependent but not sex- or tissue-dependent, however the amounts of virus-derived small RNAs varied depending on the sex and on whether or not the infected tissue was a reproductive organ.

### 3.3. *Analysis of AEFV-derived small RNAs in the co-infected* Ae. albopictus*-Vietnam*

To further characterize the small RNA profiles to AEFV (a positive-sense ssRNA virus), the 21 nt vsiRNAs and 26-30 nt vpiRNA-like small RNAs were mapped on the AEFV genome sequence. The vsiRNAs were detected across the entire viral genome in carcasses with the highest concentration of vsiRNAs mapping to the 3’ UTR (Figure 3A). The 26-30 nt small RNAs mostly mapped on the positive strand of AEFV except for the ones derived from the 3’ UTR (Figure 3B). The detected number of 26-30 nt reads in ovaries was close to background levels. Although a considerable number of 26-30 nt small RNAs reads was observed in female carcasses and male testis and carcass samples, particularly from the positive strand (Figure 2C and 3B), these small RNAs did not display the characteristic piRNA 1U or 10A bias (positions indicated by black arrows in the heat map), suggesting that they are not canonical vpiRNAs but mostly the result of degradation of viral RNA.

**Figure 3.**
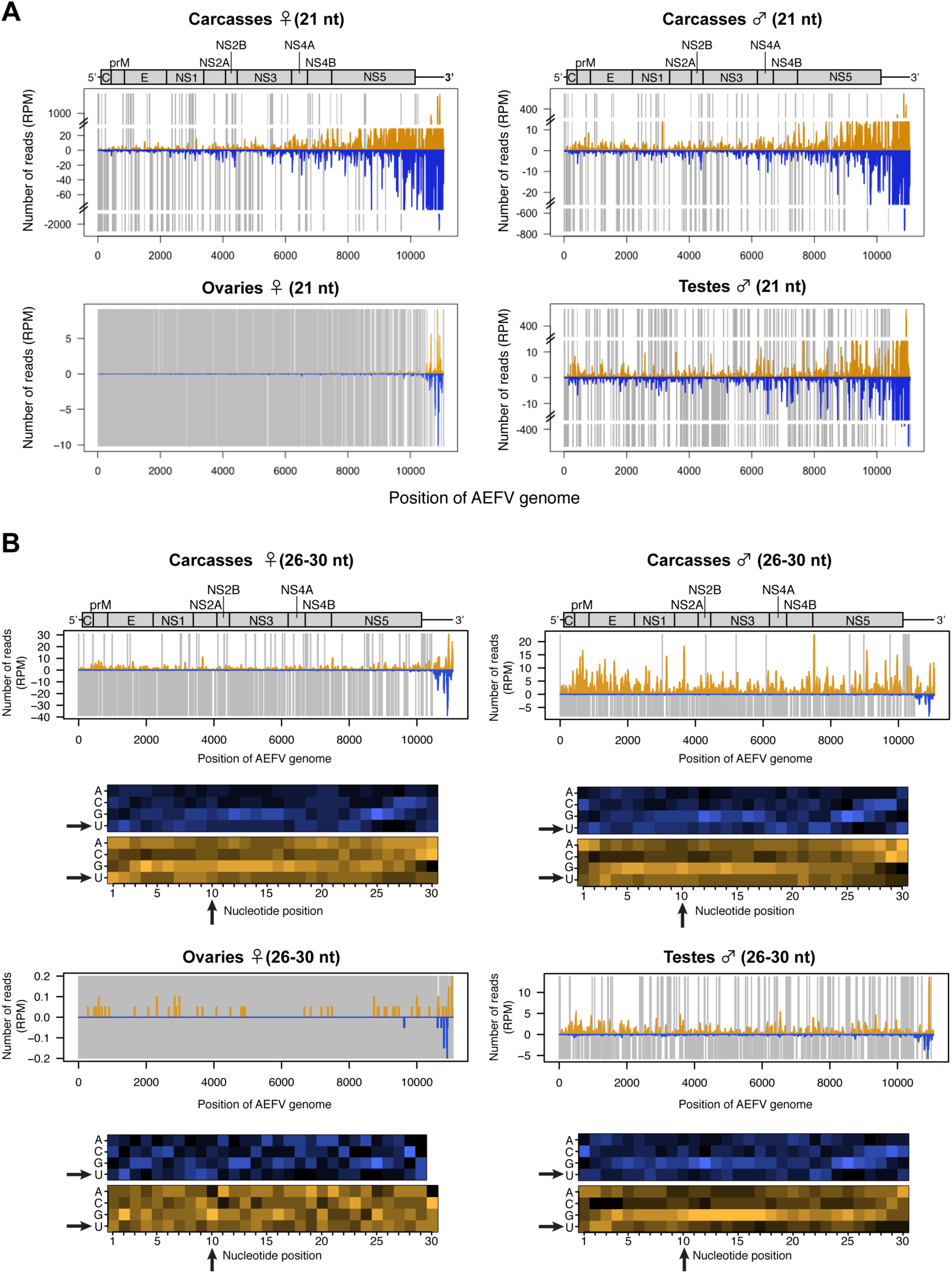
Small RNA responses to AEFV in female and male *Ae. albopictus* reproductive tissue and carcass. The distribution of siRNA (21 nt) (A) and piRNA-like small RNA (26-30 nt) (upper panel, B) mapped to AEFV genome. Schematic illustration of AEFV genome is represented on top of small RNA mapping on the viral sequence. Yellow and blue bars represent positive- and negative-stranded reads, respectively. Uncovered regions are represented as gray bars. For piRNA-like small RNA, relative nucleotide frequency per position of the 26-30 nt AEFV-derived small RNAs is shown as a heat map (lower panel in B) in which the color intensity denotes the frequency. The black arrows point to 1U and 10A positions.

### 3.4. *Analysis of SHTV -derived small RNAs in the co-infected* Ae. albopictus*-Vietnam*

Mapping of the small RNA reads on the viral genome of SHTV (a positive-sense ssRNA virus) showed that the vast majority were vsiRNAs. SHTV-derived vsiRNAs mapped across the entire genome and lacked any obvious hotspot (Figure 4A). In the ovaries, a few vsiRNAs from each position of SHTV genome (Figure 4A) and no 26-30 nt-long small RNAs (data not shown) were detected, suggesting that SHTV infection does not produce small RNAs in female germline, probably due to a lack of replication in this tissue (Figure 1). For vpiRNA production, 26-30 nt-long small RNA reads were observed in female carcasses as well as in male carcasses and testes with a random distribution through the genome (Figure 4B). No 1U or 10A biases were detected suggesting, as for AEFV, that these small RNAs are not vpiRNAs but random products of viral RNA degradation.

**Figure 4.**
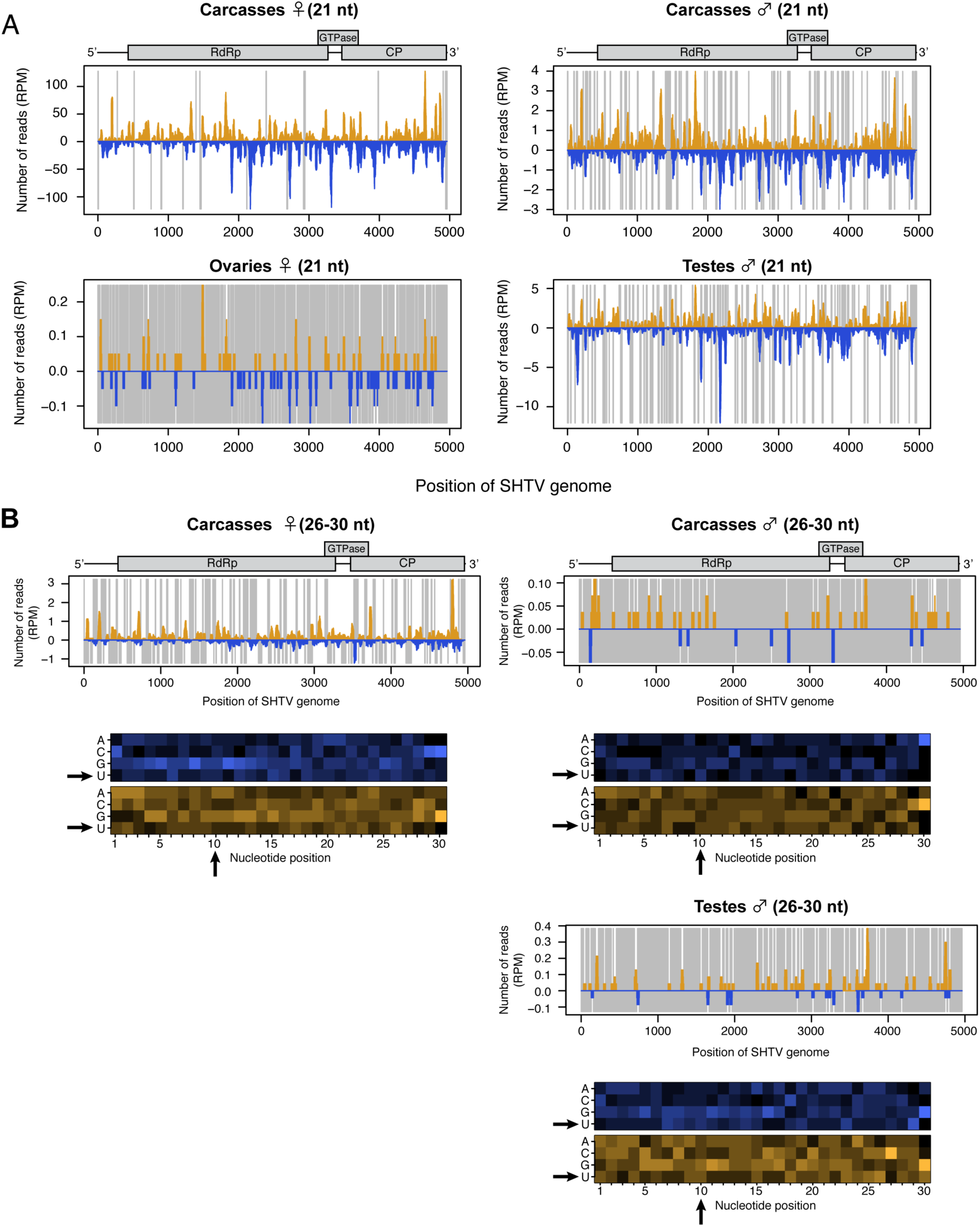
Small RNA responses to SHTV in female and male *Ae. albopictus* reproductive tissue and carcass. The distribution of siRNA (21 nt) (A) and piRNA-like small RNA (26-30 nt) (upper panel, B) mapped to SHTV genome. Schematic illustration of SHTV genome is represented on top of small RNA mapping on the viral sequence. Yellow and blue bars represent positive- and negative-stranded reads, respectively. Uncovered regions are represented as gray lines. For piRNA-like small RNA, relative nucleotide frequency per position of the 26-30 nt SHTV-derived small RNAs is shown as a heat map (lower panel in B) in which the color intensity denotes the frequency. 26-30 nt small RNA reads were not detected in ovary samples (B). The black arrows point to 1U and 10A positions.

### 3.5. *Analysis of MERV-derived small RNAs in the co-infected* Ae. albopictus*-Vietnam*

The small RNA profile of MERV (a negative-sense ssRNA virus) showed high levels of both vsiRNAs and putative vpiRNAs (Figure 2B, C and S1). We further characterized viral genomic regions producing vsiRNAs from this virus. The 21-nt vsiRNAs were distributed across the viral genome with a hotspot in the 5’ UTR region in negative-strand orientation (Figure 5A). In contrast, the 26-30-nt small RNAs mostly mapped on structural genes (N, P M and G) of the MERV genome, especially on the positive strand (Figure 5B). We observed a net 1U bias on the positive strand and 10A bias on the negative strand in all conditions (Figure 5B, red arrows), strongly suggesting that mosquitoes produced canonical MERV-vpiRNAs and induced ping-pong amplification in both reproductive tissues and carcasses. It has been reported that endogenous viral elements generate piRNAs in mosquitoes [46]. To examine whether MERV-derived piRNAs were a specific product of virus replication or derived from genomic DNA, we performed a BLASTn search for MERV-like DNA sequences in the *Ae. albopictus* genome using its genomic DNA sequence (Bioproject PRJNA587399). No MERV-like DNA sequences were found in *Ae. albopictus*-Vietnam genome. These results indicates that the vpiRNAs were effectively derived from MERV replication.

**Figure 5.**
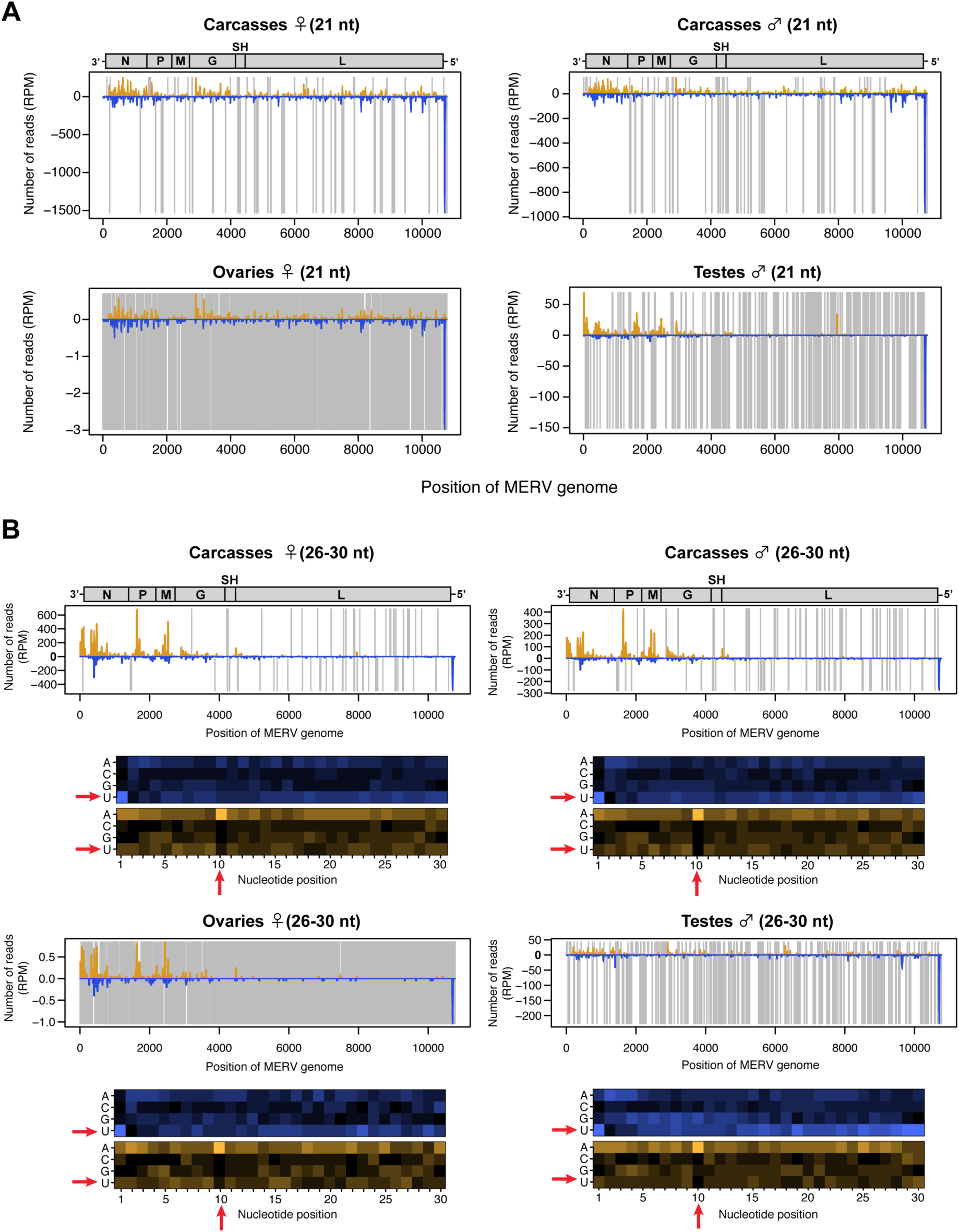
Small RNA responses to MERV in female and male *Ae. albopictus* reproductive tissue and carcass. The distribution of siRNA (21 nt) (A) and piRNA-like small RNA (26-30 nt) (upper panel, B) mapped to MERV genome. Schematic illustration of MERV genome is represented on top of small RNA mapping on the viral sequence. Yellow and blue bars represent positive- and negative-stranded reads, respectively. Uncovered regions are represented as gray lines. For piRNA-like small RNA, relative nucleotide frequency per position of the 26-30 nt MERV-derived small RNAs is shown as a heat map (lower panel in B), in which the color intensity denotes the frequency. The red arrows point to 1U and 10A positions.

### 3.6. *RNAi and PIWI genes expression in* Ae. albopictus *reproductive organs and carcasses*

A previous transcriptomic study in *Ae. aegypti* has shown differential expression of some PIWI genes between ovaries and carcasses [47]. So far, no study has analyzed or reported the function of any of the PIWI genes in *Ae. albopictus in vivo*. This prompted us to study if a link between the production of vpiRNAs and PIWI gene expression levels in co-infected mosquitoes existed. We examined mRNA levels of the main components of the piRNA pathway, together with the siRNA pathway, in ovaries, testes and carcasses. Based on *Ae. aegypti* PIWI genes, a recent bioinformatics study [35] predicted the members of the PIWI family in *Ae. albopictus* mosquitoes, Piwi1-9 and Ago3. With the same RNA samples used for the ISV RNA levels, we performed RT-qPCR for the siRNA and piRNA predicted genes. Due to the sequence similarity of some PIWI genes, we assessed gene expression as groups for Piwi1-4, Piwi5/6 and Piwi8/9. Relative gene expression was calculated by normalization with actin and with RPL18 RNA. Overall, ovaries displayed a significantly higher level of expression for all genes examined, especially PIWI genes, as compared to other samples (Figure 6A and B) implying that both small RNA pathways are present and functional in ovaries. The most remarkable difference in relative gene expression levels between reproductive tissues and carcasses was Piwi1-4. We observed that Piwi1-4 was almost exclusively expressed in ovaries and testes but not in carcasses where vpiRNAs were largely produced. Although nothing is known about a direct correlation between the expression levels of Piwi1-4 and vpiRNA production, this observation strongly suggests that Piwi1-4 is not involved in vpiRNA production in both *Ae. albopictus* female and male somatic tissues.

**Figure 6.**
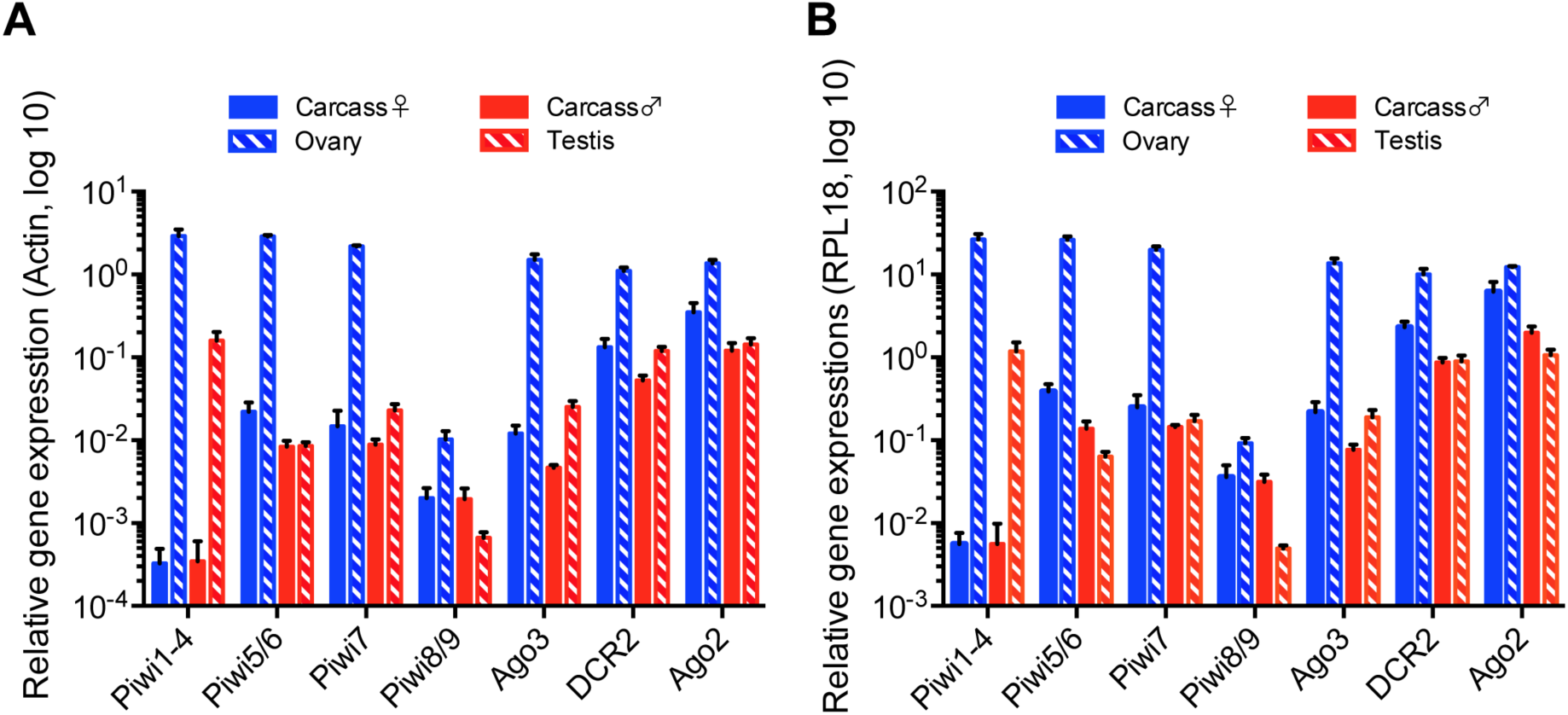
Relative gene expression of siRNA and piRNA pathway components in reproductive tissue and carcass of *Ae. albopictus* female and male co-infected with AEFV, MERV and SHTV. Relative gene expression of PIWI genes Piwi1-9 and Ago3, together with DCR2 and Ago2, key components of siRNA pathway, were examined by RT-qPCR. Relative gene expressions were calculated by normalization with actin (A) or RPL18 RNA (B). Multiple unpaired t-test with Holm-Sidak correction was applied to calculate the statistical significance. The P values are summarized in Table S2 and S3.

## 4. Discussion

Recent advances in metagenomics revealed that mosquitoes are abundantly infected with diverse ISVs in nature. However, mosquito-virus interaction studies have mainly focused on arboviruses. To better understand the small RNA responses to ISVs in mosquitoes, we used *Ae. albopictus* mosquitoes co-infected with three distant ISVs, AEFV, MERV and SHTV, as a working model. All three ISVs infect both female and male mosquitoes and are kept in the population by vertical transmission. Taking advantage of this characteristic of ISVs, we examined whether adult males mosquitoes mount the same small RNA response as adult females in their somatic tissues and reproductive tissues where the piRNA pathway is known to be active.

In the co-infected mosquitoes, all three ISVs replicated better in the carcasses compared to the reproductive tissues. SHTV replication varied among biological replicates and their RNA levels were significantly lower than AEFV and/or MERV, even if the input viral RNA copies were comparable. In *Ae. albopictus*-Japan, a significantly larger amount of SHTV-derived vsiRNAs, supporting higher levels of viral replication, was observed compared to *Ae. albopictus*-Vietnam. The lower infection level of SHTV in the Vietnam colony could be due to differences in virus susceptibility between mosquito colonies.

The overall small RNA profiles of ovaries and testes were visibly different from the rest of the body and mainly showed piRNAs rather than siRNAs or miRNAs, consistent with a previous study in *Ae. aegypti* [47]. Interestingly, testes exhibit a higher proportion of total and viral siRNAs to piRNAs when compared to ovaries. The number of siRNA reads derived from all three ISVs were significantly higher in the testes than the ovaries. This result suggests that testes might have a more active small RNA machinery or the capacity to internalize small RNAs from other virus-infected tissues and/or cells.

The production of different types of small RNAs is in agreement with previous studies on *Aedes* or *Culex* mosquito cell lines persistently infected with ISVs showing production of ISV-derived siRNAs and/or piRNAs [28, 30, 32]. While vsiRNA production was observed for all three ISVs, vpiRNAs with a clear ping-pong signature were only derived from MERV in both reproductive tissues and carcasses. Previous studies in *Aedes* and *Culex* mosquito cell lines [28, 30, 48, 49] showed that infection with Phasi Charoen-like virus (*Phenuiviriade)*, Merida virus (*Rhabdoviridae*), La Crosse virus, and Rift Valley fever virus (both *Bunyaviridae*), all of them negative-sense single-stranded viruses, produce vpiRNAs with ping-pong signature. In addition, both primary and secondary vpiRNAs were generated from some alphaviruses, positive-sense single-stranded virus, such as SINV, SFV and chikungunya virus *in vitro* and/or *in vivo* in *Aedes* mosquitoes [15, 26, 27, 48, 50]. These observations suggested that small RNA responses to ISVs are highly virus-dependent, possibly related to their replication and transcription mechanism, but independent of mosquito species or tissues.

The vsiRNAs derived from AEFV and MERV clustered in the 3’ UTR and 5’ UTR, respectively. This accumulation was also observed for 26-30 nt small RNAs although the number of reads were much lower than vsiRNAs. We speculate that these viral small RNAs from hotspots could derive from secondary structure of the virus rather than replication intermediates: DCR2 could cleave the hairpin structure of viral RNAs and the loop structure adjacent to the cleavage site might result in 26-30 nt long small RNAs as by-product of vsiRNAs production. Interestingly, previous studies suggested that the subgenomic RNA of flaviviruses, which corresponds to the 3’ UTR, is capable of supressing the RNAi machinery, possibly as an RNA decoy or by inhibiting the loading of vsiRNA into the RISC [21, 23, 24]. The high amount of siRNAs from the the 3’ UTR of AEFV could be a strategy from the virus to escape from the siRNA pathway. Further analyses are required to explore this hypothesis.

MERV-derived piRNAs were largely produced from the viral genomic region of structural genes corresponding to nucleotide positions 130 to 4162. *Cx. quinquefasciatus* cell lines infected with Merida virus, also a rhabdovirus, showed a similar vpiRNA profile, suggesting that the regions that generate vpiRNAs are conserved regardless of the mosquito species and depend on the viral replication strategy [30]. Rhabdoviruses, in general, transcribe their mRNA from the 3’ UTR, which is closer to structural genes. The structural gene transcripts can be more efficiently targeted by the piRNA pathway compared to the non-structual genes located in the 5’ end. Interestingly, different studies showed that vpiRNAs from alphaviruses, orthobunyaviruses and phelobovirus were also largely generated from viral genomic regions encoding structural genes *in vitro* and/or in *Aedes* mosquitoes [15, 26, 27, 48-50]. It remains to be elucidated how exogenous viral RNAs are recognized as precursors for vpiRNA biogenesis, however, transcription mechanism of viral structural genes might be involved in recognition by PIWI proteins. In *Ae. aegypti*, Piwi5 is likely to play a role in producing primary piRNAs from both TEs and exogenous viruses, however how Piwi5 selects the transcript to process into primary piRNAs is still unknown [26].

A recent study identified ten PIWI genes, Piwi1-9 and Ago3, in *Ae. albopictus* based on PIWI genes in *Ae. aegypti* using tBLASTx search [35]. However, it is important to note that the genome assembly and gene annotation of *Ae. albopictus* is not as well developed as compared to *Ae. aegypti*. We quantified the expression levels of the putative Piwi1-9 and Ago3 together with DCR2 and Ago2 genes by qPCR in ISVs-infected *Ae. albopictus* samples. A previous study examined PIWI gene expression in *Ae. albopictus* female midguts and the rest of bodies by transcriptome analysis and RT-PCR [35]. Our results were consistent with this study, for instance Piwi1-4 were abundantly expressed in the reproductive tissues, which correspond to the non-midgut body samples of the transcriptome analysis. Our qPCR analysis detected Piwi8/9 expression in both adult reproductive tissues and the carcasses, while their expression was only observed in embryo and larvae stage by RT-PCR in the previous study. This difference could be explained by the sensitivity of the assay. Interestingly, our study showed that Piwi1-4 were exclusively expressed in the reproductive tissues that has significantly less MERV-derived vpiRNAs than the carcass. This observation indicates that Piwi1-4 do not play a role on vpiRNA production; in other words, vpiRNA biogenesis requires one or some of Piwi5-9 in *Ae. albopictus*. Nevertheless, a better genome assembly and annotation for *Ae. albopictus* is required to conduct studies that will allow to determine the PIWI genes involved in vpiRNA biogenesis.

The present study demonstrated that, regardless of the mosquito sex and the reproductive nature of the tissue, production of different types of viral small RNAs from ISVs are virus-dependent. Our results highlight the value of ISVs as virus models to better understand small RNA pathways in mosquito as well as the potential countermeasures of viruses to escape the RNA interference response. Future studies should address the small RNA responses to ISVs in mosquitoes that are naturally infected. This could lead to new insights into mosquito immunity and mosquito – ISV – arbovirus relationships.

## Supplementary Materials

Figure S1: Small RNA size distribution mapped to each ISV in reproductive tissues and carcasses of female or male *Ae. albopictus* mosquitoes, Figure S2: Small RNA profile of SHTV in reproductive tissues and carcasses of female *Ae. albopictus*-Japan mosquitoes, Table S1: Primer sequences used, Table S2: P values for qPCR analysis normalized with actin in Figure 6, Table S3: P values for qPCR analysis normalized with RPL18 in Figure 6.

## Author Contributions

Conceptualization, L.F., M.C.S. and Y.S.; methodology, L.F., H.B., and Y.S.; investigation, L.F., H.B. and Y.S.; resources, M.C.S.; formal analysis, L.F. and Y.S.; writing, —original draft preparation, Y.S.; writing—review and editing, all authors.

## Acknowledgments

We thank all members of the Saleh lab for discussion and Susan Carpenter and Louis Lambrechts for critical reading and editing of the manuscript, Louis Lambrechts for providing mosquito colonies, Annabelle Henrion-Lacritick and Catherine Lallemand for assistance with mosquito rearing, Haruhiko Isawa for providing the AEFV/MERV/SHTV stock and mosquito colonies.

