## Supplemental Material for "Differential small RNA responses against co-infecting insect-specific viruses in *Aedes albopictus* mosquitoes"

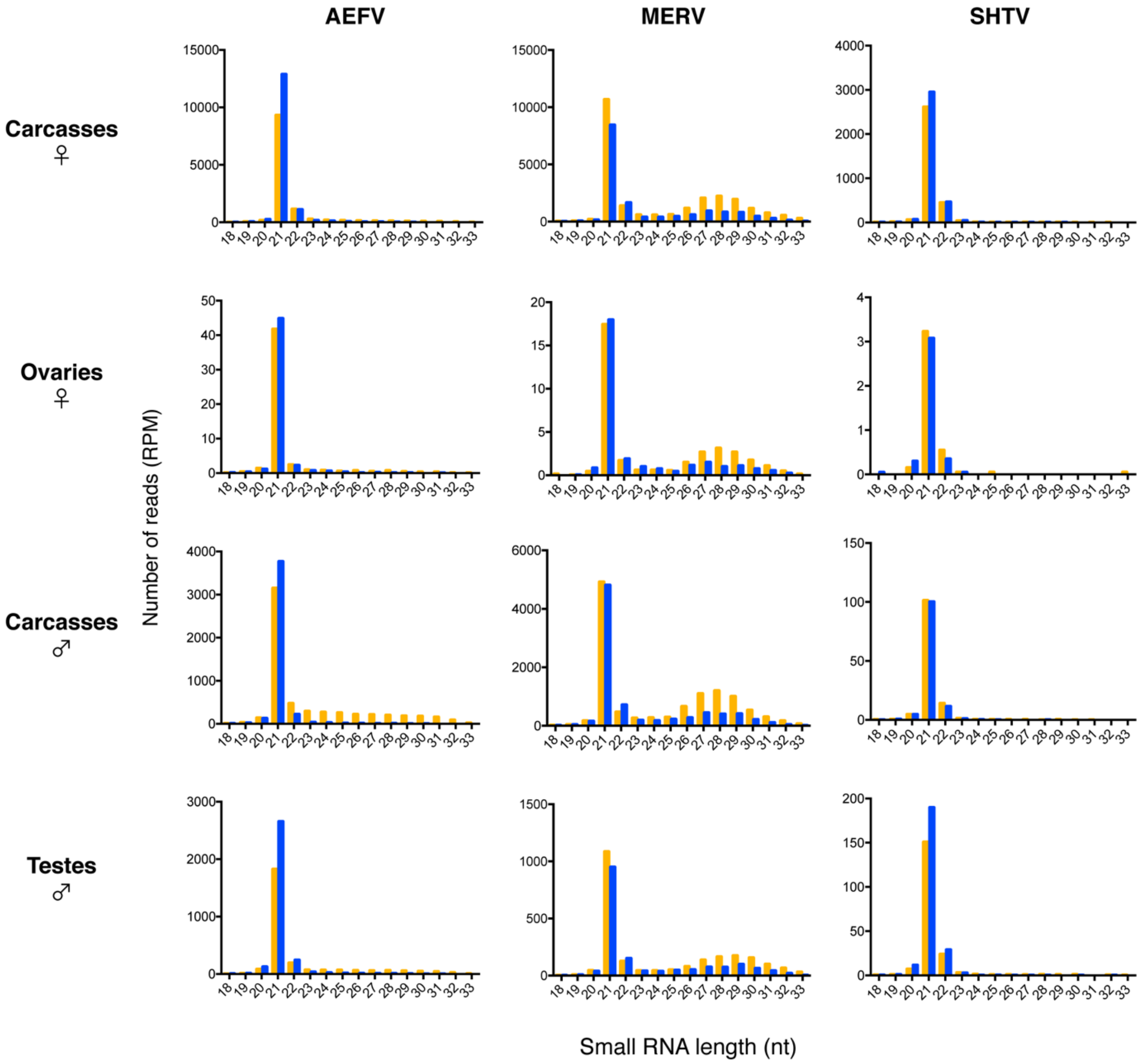


**Figure S1.** Small RNA size distribution mapped to ISVs in reproductive tissues and carcasses of female or male *Ae. albopictus* mosquitoes. Yellow and blue bars represent positive- and negative-stranded reads, respectively.

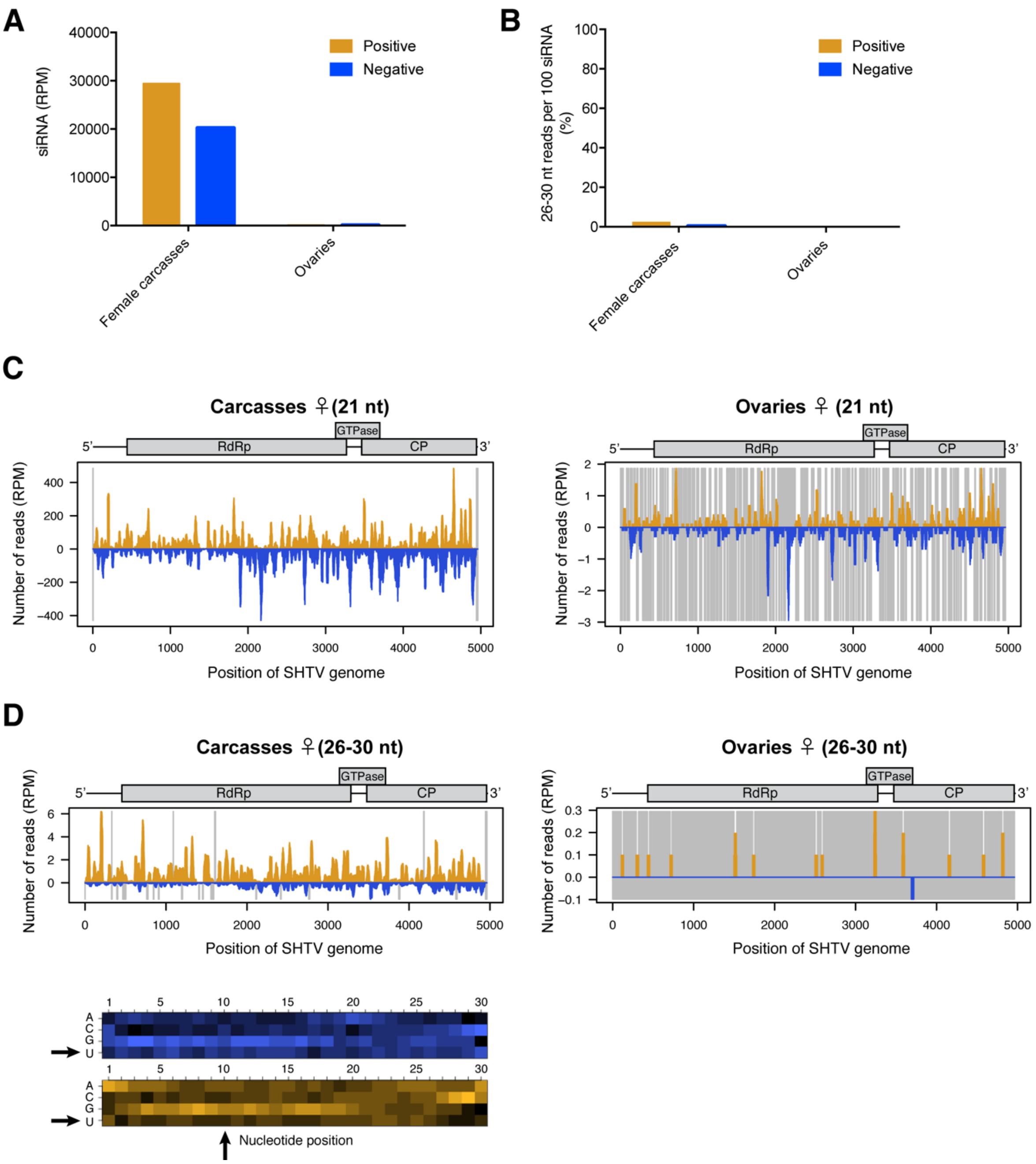


**Figure S2.** Small RNA profile of SHTV in reproductive tissues and carcasses of female *Ae. albopictus*-Japan mosquitoes. Normalized siRNA reads per one million (RPM) mapped on SHTV genome is shown in (A). Proportion of 26-30 nt long SHTV-derived small RNAs to 100 reads of its vsiRNAs is shown in (B). The distribution of siRNA (21 nt) (C) and piRNA-like small RNA (26-30 nt) (upper panel, D) mapped to SHTV genome is shown. Yellow and blue bars represent positive- and negative-stranded reads, respectively. Uncovered regions are represented as gray lines. For piRNA-like small RNA, relative nucleotide frequency per position of the 26-30 nt SHTV-derived small RNAs in carcasses is shown as a heat map (lower panel in D), in which the color intensity denotes the frequency. 26-30 nt small RNA reads were not detected in ovary samples (B). The black arrows point to 1U and 10A positions.

Table S1. Primer sequences used.

| Purpose | Sequences |
|---|---|
| AEFV_NS4 RNA standard | Forward: GGATCCTAATACGACTCACTATAGGAGTTCAGCTAGTCTGTACGACATCC |
| | Reverse: GCGAAATTTACTGAAAGGGGCCATAG |
| qPCR for AEFV | Forward: AGGCACAGTTGGAGGGTTCC |
| | Reverse: TACAACGCTGGGAAGGCCAA |
| qPCR for MERV | Forward: GAGGTGTCGGATAAATTTCTAG |
| | Reverse: TTATCTGATAGGTGCCCCTTC |
| qPCR for SHTV | Forward: TTCTTAGGAATGAGGCTCATG |
| | Reverse: ACTAGGTCGTTGGGGCTCTC |
| qPCR for DCR2 | Forward: CCAAAACCGCTGAAGGAGA |
| | Reverse: CTGGACAATCAATCCGAGCA |
| qPCR for Ago2 | Forward: GTCGGGTGGATTTGGTGTTC |
| | Reverse: TATCGCCCCGTTTCCTATCC |
| qPCR for Piwi1-4 | Forward: CGACACGAACGACAAATCCA |
| | Reverse: GGTACTCGTTGAGCGCCTTG |
| qPCR for Piwi5/6 | Forward: CGCAGTTGGTGATGTGTGTG |
| | Reverse: ATGACTTGCGTGGGAATGG |
| qPCR for Piwi7 | Forward: CTGAAGACCCGAACGATCAC |
| | Reverse: TTACCATCACCGAAGCCAGA |
| qPCR for Piwi8/9 | Forward: GCAGTGGTTCGCAGTGGTTC |
| | Reverse: CCGGCGAATCGTTGGGAATG |
| qPCR for Ago3 | Forward: CCATTCGCCGGACATTCTGC |
| | Reverse: TACTGACAGCAAGCGGGGAC |

Table S2. P values for qPCR analysis normalized with actin in Figure 6

| | **Carcass_♀ vs Ovaries** | **Carcass_♂ vs Testes** | **Carcass_♀ vs Carcass_♂** | **Ovaries vs Testes** | **Ovaries vs Carcass_♂** | **Carcass_♀ vs Testes** |
|---|---|---|---|---|---|---|
| DCR2 | 1,49E-04 | 1,98E-03 | 1,85E-02 | 1,23E-04 | < 0,0001 | 6,05E-01 |
| Ago2 | 5,90E-04 | 3,61E-01 | 1,94E-02 | 1,41E-04 | 1,33E-04 | 2,66E-02 |
| Piwi1-4 | 9,71E-04 | 3,16E-03 | 9,15E-01 | 1,22E-03 | 9,71E-04 | 3,16E-03 |
| Piwi5/6 | < 0,0001 | 8,86E-01 | 2,46E-02 | < 0,0001 | < 0,0001 | 2,43E-02 |
| Piwi7 | < 0,0001 | 5,20E-03 | 2,81E-01 | < 0,0001 | < 0,0001 | 1,89E-01 |
| Piwi8/9 | 6,06E-03 | 2,91E-02 | 9,41E-01 | 3,15E-03 | 5,96E-03 | 2,61E-02 |
| Ago3 | 4,96E-04 | 1,30E-03 | 1,22E-02 | 5,14E-04 | 4,87E-04 | 1,29E-02 |

Table S3. P values for qPCR analysis normalized with RPL18 in Figure 6

| | **Carcass_♀ vs Ovaries** | **Carcass_♂ vs Testes** | **Carcass_♀ vs Carcass_♂** | **Ovaries vs Testes** | **Ovaries vs Carcass_♂** | **Carcass_♀ vs Testes** |
|---|---|---|---|---|---|---|
| DCR2 | 1,32E-03 | 8,23E-01 | 3,37E-03 | 6,32E-04 | 6,19E-04 | 4,09E-03 |
| Ago2 | 5,74E-03 | 1,88E-02 | 1,94E-02 | < 0,0001 | < 0,0001 | 9,55E-03 |
| Piwi1-4 | 5,75E-04 | 3,52E-03 | 9,99E-01 | 6,93E-04 | 5,75E-04 | 3,52E-03 |
| Piwi5/6 | < 0,0001 | 1,58E-02 | 9,03E-03 | < 0,0001 | < 0,0001 | 2,89E-03 |
| Piwi7 | < 0,0001 | 2,46E-01 | 1,38E-01 | < 0,0001 | < 0,0001 | 2,50E-01 |
| Piwi8/9 | 8,92E-03 | 2,76E-03 | 6,58E-01 | 5,31E-04 | 3,13E-03 | 1,85E-02 |
| Ago3 | 3,06E-04 | 1,15E-02 | 2,63E-02 | 3,03E-04 | 2,93E-04 | 5,71E-01 |
